# Adult mouse retina explants: an ex vivo window to explore central nervous system diseases

**DOI:** 10.1101/2020.02.22.960609

**Authors:** Julia Schaeffer, Celine Tardy, Floriane Albert, Stephane Belin, Homaira Nawabi

## Abstract

When the developing central nervous system (CNS) becomes mature, it loses its ability to regenerate. Therefore, any insult to adult CNS leads to a permanent and irreversible loss of motor and cognitive functions. For a long time, much effort has been deployed to uncover mechanisms of axon regeneration in the CNS. It is now well understood that neurons themselves lose axon regeneration capabilities during development, and also after a lesion or in pathological conditions. Since then, many molecular pathways such as mTOR and JAK/STAT have been associated with axon regeneration. However, no functional recovery has been achieved yet. Today, there is a need not only to identify new molecules implicated in adult CNS axon regeneration, but also to decipher the fine molecular mechanisms associated with regeneration failure. This is critical to make progress in our understanding of neuroprotection and neuroregeneration and for the development of new therapeutic strategies. In this context, it remains particularly challenging to address molecular mechanisms in in vivo models of CNS regeneration. The extensive use of embryonic neurons as in vitro model is a source of bias, as they have the intrinsic competence to grow their axon upon injury, unlike mature neurons. In addition, this type of dissociated neuronal cultures lack a tissue environment to recapitulate properly molecular and cellular events in vitro. Here, we propose to use cultures of adult retina explants to fill this gap. The visual system - which includes the retina and optic nerve - is a gold-standard model to study axon regeneration and degeneration in the mature CNS. Cultures of adult retina explants combine two advantages: they have the simplicity of embryonic neurons cultures and they recapitulate all the aspects of in vivo features in the tissue. Importantly, it is the most appropriate tool to date to isolate molecular and cellular events of axon regeneration and degeneration of the adult CNS in a dish. This ex vivo system allows to set up a large range of experiments to decipher the fine molecular and cellular regulations underlying mature CNS axon growth.

## INTRODUCTION

For decades, many efforts have been deployed to unlock the cellular programs to achieve axon regeneration in the central nervous system (CNS). Indeed, unlike neurons from the peripheral nervous system (PNS), CNS neurons are not able to grow axons after injury. The pioneering work from David and Agayo(*1*) highlighted that environment modulation of lesioned CNS axons could, to some extent, help axonal growth. Those studies opened up new venues to address CNS regeneration mechanisms. Different contributors of the axonal environment were subsequently identified to impair axon regeneration. After CNS injury, axons are demyelinated and the release of myelin debris (composed of growth-inhibitory molecules such as Nogo, MAG or OMpg) in their surroundings contributes to inhibit any growth attempt(*2*). In addition, a glial scar, which confines locally the inflammation process that occurs following the lesion, forms at the injury site. This scar acts as a physical barrier and is also a source of inhibitory molecules such as chondroitin sulfate protoglycans (CSPG) or repulsive guidance molecules(*3*). Using embryonic neuron cultures, many studies deciphered the molecular and cellular mechanisms underlying their inhibitory role on neurite outgrowth. However, in vivo experiments in mouse models of CNS injury revealed that those mechanisms account only partially for the regeneration failure of the CNS(*4*). Therefore, other hypotheses emerged and focused on neurons themselves(*5*). It is now well understood that CNS regeneration aborts because of a dual mechanism: on one hand neurons lose the ability to grow during development and on the other hand, the injury itself inhibits further axon regrowth(*5, 6*). Therefore, the activation of developmentally regulated pathways such as mTOR (mammalian target of rapamycin)(*5*), which is a master regulator of protein translation and cell growth, or transcription factors such as KLF (kruppel like factors)(*7*) promotes axon regeneration in different models of CNS injury, such as optic nerve or spinal cord injuries(*5, 7–9*). Moreover, as the lesion itself modulates several signaling pathways, their synergistic manipulation promotes long distance regeneration(*6, 10*).

Even though many exciting candidates regulating axon regeneration have been uncovered, their modulation in a therapeutic approach remains difficult. Indeed, most of these molecules trigger numerous functions in cells and current knowledge is insufficient to understand which one is essential for axon regeneration. In addition, these regenerative molecules are also known to be oncogenic factors(*11*). Therefore, it is urgent to unravel the precise molecular and cellular events allowing axon regeneration and elongation in mature CNS in order to i) characterize new cellular targets implicated in axon regeneration mechanisms, and ii) develop innovative therapeutic strategies for CNS repair after a traumatic lesion or in neurodegenerative diseases. In this regard, embryonic cortical or hippocampal neuronal cultures are commonly used as experimental models(*8, 12*). However, while it is easy to obtain a large quantity of isolated neurons in culture, these in vitro models cannot answer fully and precisely to the question of axon regeneration in the mature CNS. Indeed, embryonic neurons and adult neurons have different intrinsic abilities when it comes to axon growth after a traumatic lesion. Embryonic neurons or young neurons (until P5-P6 in mice) have great regrowth potential(*13, 14*). On the contrary, this feature is lost in mature neurons. For this reason, it is crucial to find in vitro assays that recapitulate in vivo features of mature neurons.

The optic nerve has proved to be a good model to address the molecular and cellular mechanisms of CNS axon regeneration in adult. Most of the molecular pathways that have been uncovered using optic nerve lesion as a model of CNS injury have also shown promising results for regeneration in the cortico-spinal tract, the main axon bundle that controls voluntary movements in humans(*9, 15*). Within the retina, only one population of neurons project their axons to form the optic nerve: the retinal ganglion cells (RGC). These neurons build the connection between the eye and the brain through the optic nerve. Unlike spinal cord lesions that affect multiple neuronal populations, the optic nerve injury affects only the population of RGC. This unique feature allows to focus specifically on the specific behavior of this population of neurons upon axon injury(*6*). Therefore, the optic nerve and RGC are one of the best models to address CNS regeneration modalities.

Here, we propose a method to translate the in vivo phenotype into an ex vivo approach to decipher the molecular and cellular mechanisms underlying mature CNS regeneration. To this aim, we use adult mouse retina explant in culture. This technique combines the simplicity of embryonic neuronal cultures, and all the characteristics of an adult system. Indeed, we show that our model recapitulates all the features observed in vivo in RGC after optic nerve injury: axon regeneration induced by intrinsic factors, growth extent, number of regenerative axons. In addition, in our set up, RGC neurons are not in dissociated in culture but are kept in a whole retinal structure. Finally, we could perform the lesion of a single axon using laser guided ablation. Similarly to embryonic neuronal cultures, the adult retina explant system is an evolutive toolbox to test several cellular functions and to study the fine mechanisms of axon growth (**Figure 1**). Treatment in culture media can be easily achieved in order to test different drugs that could potentiate axon growth or block cellular pathways. These cultures allowed us also to study growth cone behavior, axon guidance modalities in adult and organelle or cytoskeleton dynamics. Importantly, we explored fine cellular mechanisms linked to mature CNS regeneration at a single axon level using this set-up. The list of experiments is not exhaustive and our system can be implemented into many experimental designs to respond to future questioning in the field. Our model is a great tool to address all the current questions regarding physiological events that are difficult to achieve in vivo.

**Figure 1:**
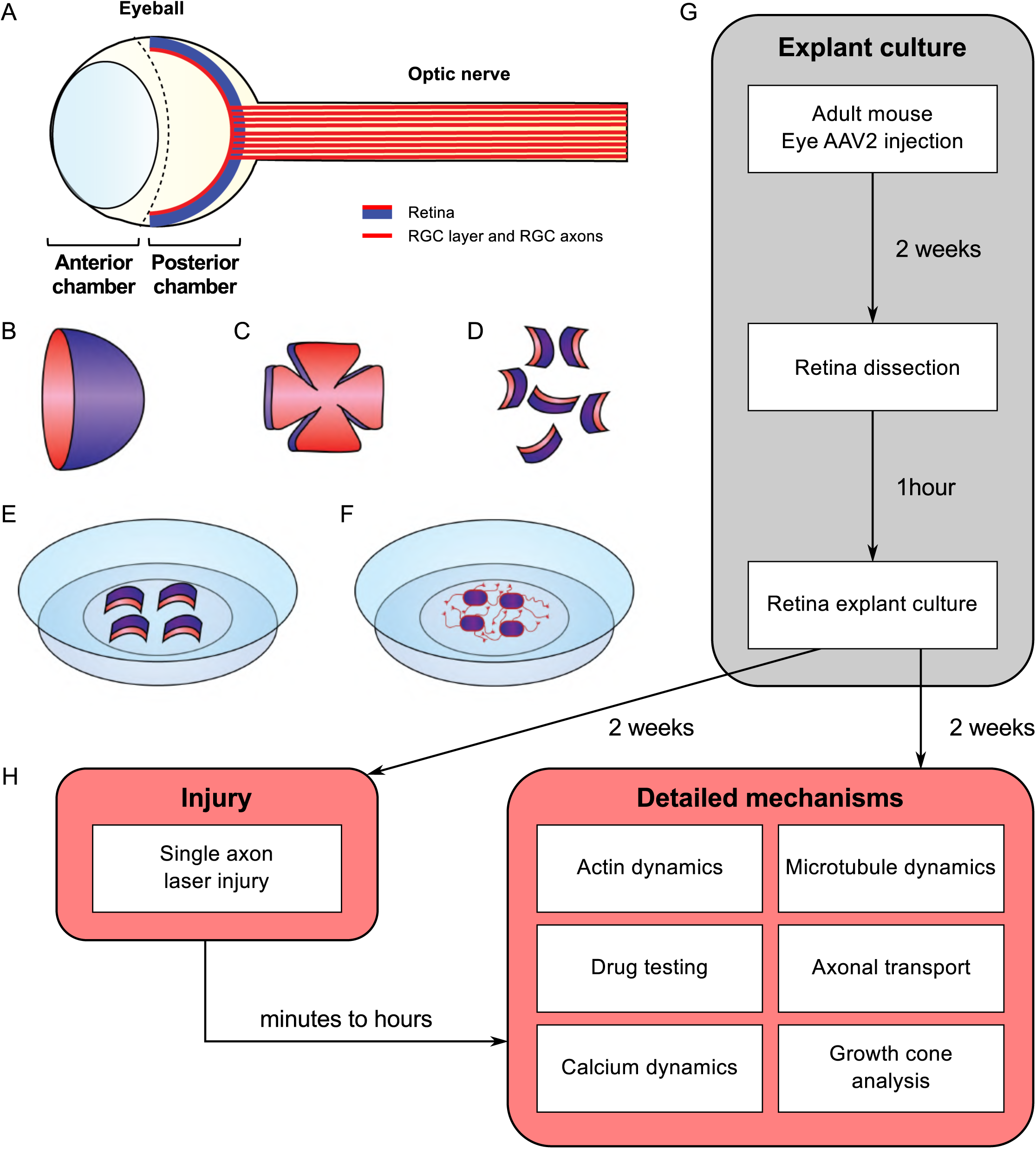
Schematic of experimental procedure for adult retina explant cultures and examples of ex vivo analyses. (A) Schematic of the eye. The retina is covering the inner side of the eye ball posterior chamber. Retinal ganglion cells are located in the anterior face of the retina (in red) and sending their axons to form the optic nerve (B) The retina dissected out of the eye ball is cup-shaped. The RGC layer (in red) is located in the inner side of the cup. (C) The retina is cut in flower shape. (D) The retina is chopped into small pieces (500µm in diameter). All the pieces will show a curvature. The RGC layer (in red) is on the concave side. (E) Put the RGC side (in red) in the coverslip coated with laminin and poly-L-lysin and covered with a thin layer of adhesion media. (F) Schematic of an adult retina explant culture with axons growing on the substrate from RGC. (G) Timeline of standard experiment. (H) Examples of downstream applications, including ex vivo single axon laser injury and/or ex vivo analysis of axon biology (e.g. cytoskeleton dynamics, growth cone dynamics, axonal transport).

## RESULTS

### Neurites are axons from RGC

The mechanisms underlying axon regeneration in mature CNS remain difficult to address. While it is common to use embryonic neuronal cultures to address those questions, the intrinsic ability of young neurons to regrow their axon induces a bias regarding the molecular and cellular mechanisms of axon growth in mature neurons(*13*). Thus, there is a need to find better ex vivo systems to decipher mature CNS regeneration processes. Adult retina and optic nerve are gold-standard models to study neuron survival and axon regeneration in the CNS. The retina is a highly organized multi-layered organ composed of 5 types of neurons: photoreceptors, horizontal cells, bipolar neurons, amacrine cells and retinal ganglion cells (RGC). RGC are the only neuronal population sending their axon to form the optic nerve, which is the only bridge between the eye and the brain. Therefore, a lesion to the optic nerve will affect specifically these neurons within the retina. Here, we managed to culture retina explants from adult mouse (**Figure 1**). Because of the intrinsic growth incompetence of adult CNS neurons, we observed that wild-type (WT) explants grow few neurites **(Table 1)**. Therefore, we activated the mTOR pathway, known to induce axon regeneration in mature CNS(*5*), through deletion of PTEN. Using the PTEN^fl/fl^/YFP-17 mouse line(*16*), we injected 1µL of AAV2-Cre into the vitreous body of the eye to delete PTEN in RGC. YFP is expressed by 99% of RGC and few amacrine cells (**Figure 2A**). Two weeks after injection, we dissected out the retinas and put the adult explants in culture (**Figure 1**). After two weeks in culture, we performed immunocytochemistry using an anti-β Tubulin III (TUJ1) antibody to label neurites. We found that 90.6% of neurites were YFP^+^, meaning that almost all the neurites that grow out of the explant tissue are from RGC (**Figure 2B**). To address whether these neurites are axons, we used an anti-Tau antibody, a specific axon marker. We found that 98.3% of YFP^+^ neurites are Tau^+^, meaning that most of the neurites growing out of the explants are RGC axons (**Figure 2C**). Therefore, this ex vivo set up is ideal to study molecular and cellular mechanisms of axon regeneration specifically in mature RGC at a single axon level.

**Table 1:** NeuriteJ quantification of axon growth on adult retina explants.

| Condition | Total number explants | Number explants with no axon growth | Number explants for NeuriteJ quantification<sup>1,2</sup> |
| --- | --- | --- | --- |
| Pten<sup>fl/fl</sup> + AAV2-Plap | 47 | 18 (38.3%) | 29 (61,7%) |
| Pten<sup>fl/fl</sup> + AAV2-Cre | 41 | 8 (19.5%) | 33 (80,5%) |
| Pten<sup>fl/fl</sup>/SOCS3<sup>fl/fl</sup> + AAV2-Cre + AAV2-CNTF + AAV2-c-myc | 64 | 6 (9.4%) | 22 (90,6%) |

Data collected from two independent days of explant culture.
<sup>1</sup>Explants with no axon growth were excluded from quantification.
<sup>2</sup>Explants with overlap of axons from neighbouring explants were excluded from quantification.

**Figure 2:**
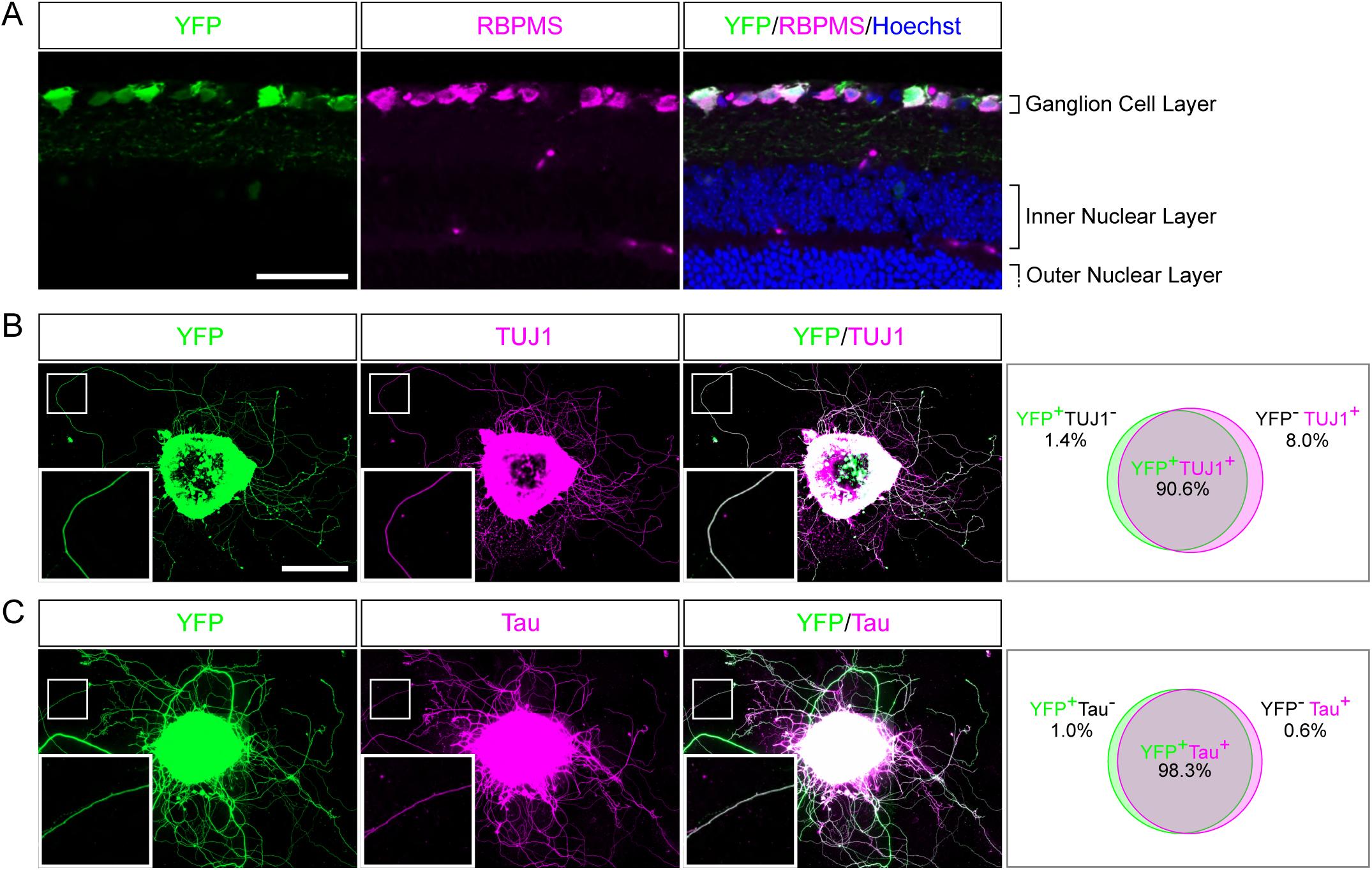
Characterization of neurites in adult retina explant culture. (A) Representative confocal picture of a PTEN^fl/fl^/Thy1-YFP adult mouse retina section. RGC expressing YFP (labelled in green) are stained with anti-RBPMS (magenta) antibody and nuclei are labelled with Hoechst (blue). Scale bar: 50µm. (B-C) Representative pictures of an adult retina explant from a PTEN^fl/fl^/Thy1-YFP mouse with priori intravitreal injection of AAV2-Cre. RGC neurites are labelled with YFP (in green), anti-β Tubulin III (TUJ1) (magenta) (B) or anti-Tau (magenta) (C), with corresponding quantification of colocalization. At least 350 neurites from at least 18 explants and at least 3 independent experiments were quantified. Scale bar: 500µm.

### Adult retina explant cultures recapitulate in vivo phenotypes

An ex vivo system is more relevant when it recapitulates the phenotype of the corresponding in vivo model. It is now largely described that wild-type adult CNS neurons are not able to grow axons after injury and that modulating neurons themselves enable CNS axon regeneration(*17*). Extensive axon regeneration in the mature CNS was achieved for the first time through the activation of the mTOR pathway(*5*). Upon its activation, axons are able to grow several hundreds of micrometers from the injury site. Since then, long distance regeneration from the eye ball to the brain has been obtained by the synergistic activation of mTOR, JAK/STAT and c-myc pathways in RGC(*6*). Therefore, we compared axon regeneration in these conditions in vivo in the optic nerve and ex vivo in adult retina explants (**Table 1 and Figure 3A**). To do so, we used PTEN-floxed (PTEN^fl/fl^) and PTEN^fl/fl^/SOCS3^fl/fl^ mouse lines, as PTEN and SOCS3 are negative regulators of mTOR and JAK/STAT pathways respectively. We injected 1µL of AAV2-Cre into the vitreous bodies of P28 PTEN^fl/fl^ to delete specifically PTEN from RGC. AAV2-Cre, AAV2-CNTF (to activate JAK/STAT pathway) and AAV2-c-myc were injected into the vitreous bodies of P28 PTEN^fl/fl^/SOCS3^fl/fl^ mice. As control, we used PTEN^fl/fl^ mice injected with AAV2-Plap. Two weeks after injection, we proceeded with optic nerve injury or with retina explant culture (**Figure 3A, B, C**). With this experimental design, we could focus on the same tissue at the same stage for in vivo and ex vivo experiments. After two weeks in culture, we analyzed axon growth in vivo and ex vivo. For the in vivo part, we injected Alexa555-conjugated cholera toxin B (CTB-555) in the vitreous body 48 hours before sacrificing the animals to label regenerative fibers. Optic nerves were subsequently dissected and cleared (**Figure 3B**), then the whole tissue was imaged with confocal microscopy to assess the extent of axon regeneration. We found that in control conditions, very few axons extended from the injury site and grew only few micrometers (**Figure 3D, E, F**). PTEN-deleted axons showed better growth with axons reaching up to 500µm from the injury site (**Figure 3D, E, F**). Strikingly, PTEN^−/−^/SOCS3^−/−^+c-myc axons are able to grow over long distances. This regeneration is very robust as axons could reach up to 1500µm from the injury site two weeks post-injury (**Figure 3D, E, F**; **Suppl Figure 1A, B, C**). The advantage of using tissue clearing is to appreciate the whole extent of regeneration and to avoid any bias induced by tissue sectioning. For quantification, manual axon counting is not accurate enough, especially close to the lesion site where axon number is high in regenerative conditions. Therefore, we set up a semi-automated method using Image J software (**Suppl Figure 1D**) to quantify the extent of regeneration without any bias. We measured CTB-555 intensity along the optic nerve at defined distances from the injury site. CTB-555 intensity was then normalized by the width of the optic nerve at each distance and by the maximal CTB-555 intensity in the regenerating region to account for experimental variability. Finally, we subtracted the background value. The resulting normalized integrated intensity at each distance allowed us to quantify the extent of axon regeneration accurately and without any bias.

**Figure 3:**
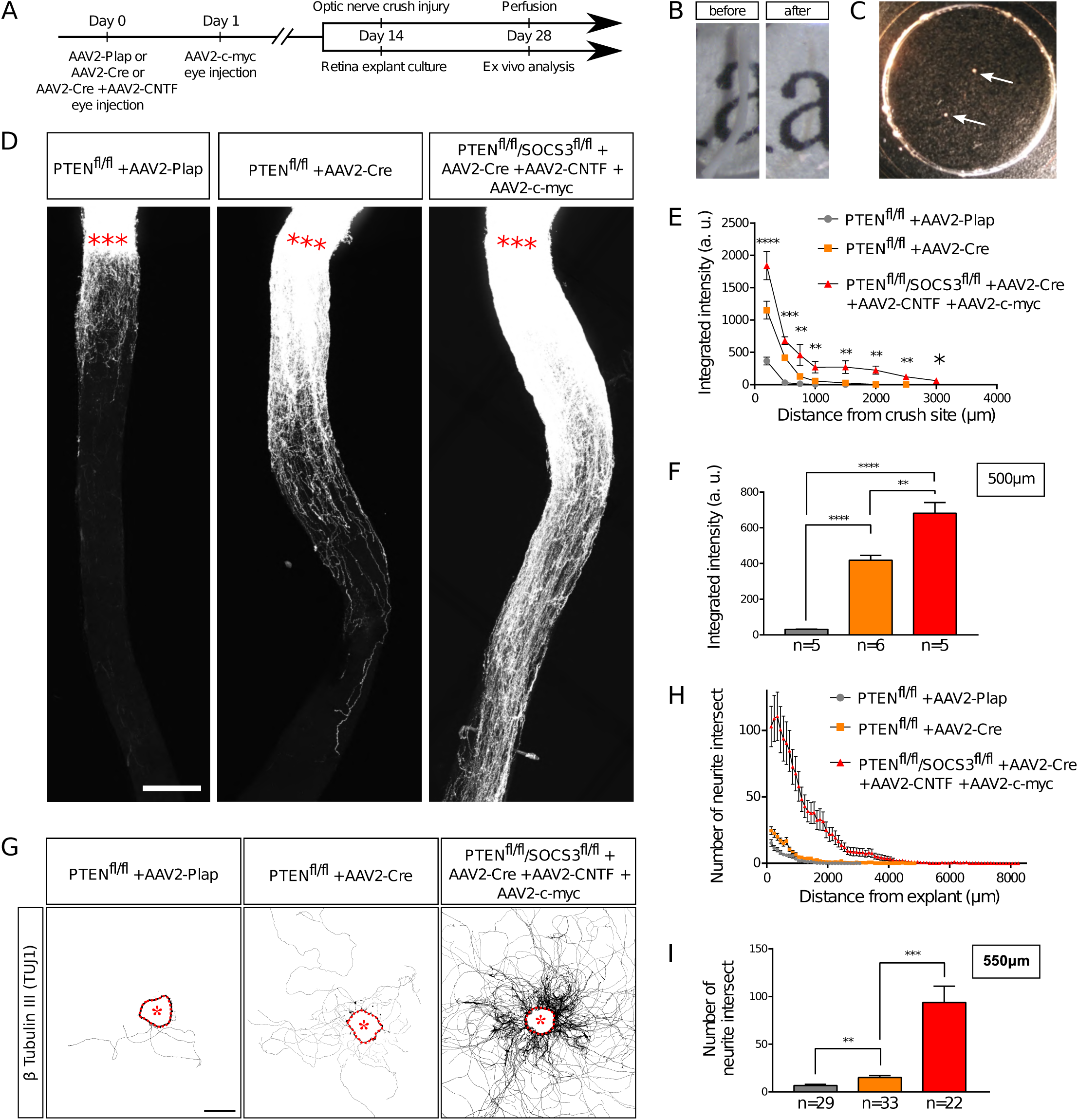
Adult retina explant culture mimics in vivo axon regeneration. (A) Timeline of experiments. (B) Picture of mouse optic nerve before and after transparisation. (C) Picture of retina explants (arrows) in culture in a glass-bottom dish. (D) Representative confocal pictures of whole optic nerve showing axon regeneration 2 weeks post-injury in control (PTEN^fl/fl^ + AAV2-Plap), PTEN-deleted (PTEN^fl/fl^ + AAV2-Cre) and PTEN/SOCS3 co-deleted c-myc-overexpressing (PTEN^fl/fl^/SOCS3^fl/fl^ + AAV2-Cre + AAV2-CNTF + AAV2-c-myc) conditions. Axons are traced with fluorescently-labelled CTB. Red stars indicate the injury site. Scale bar: 200µm. (E) Quantification of axon regeneration (integrated intensity) in (D). Data expressed as mean +/- s.e.m. ANOVA test. (F) Quantification of axon regeneration at 500µm of injury site. Unpaired t-tests. * p<0.05, ** p<0.01, *** p<0.001, **** p<0.0001. (G) Representative pictures of adult retina explants after 2 weeks in culture, from control (PTEN^fl/fl^ + AAV2-Plap), PTEN-deleted (PTEN^fl/fl^ + AAV2-Cre) and PTEN/SOCS3 co-deleted + c-myc-overexpressing (PTEN^fl/fl^/SOCS3^fl/fl^ + AAV2-Cre + AAV2-CNTF + AAV2-c-myc) conditions. Axons are labelled with anti-β Tubulin III (TUJ1) antibody. The star indicates the explant. Scale bar: 500µm. (H) Quantification of axon growth in (G) with Scholl analysis. Data expressed as mean +/- s.e.m. (I) Quantification of axon growth (number of axon intersect) at 550µm of explant. Mann-Whitney tests. ** p<0.01, *** p<0.001.

In parallel, after two weeks in culture, explants were fixed and stained with an anti-β Tubulin III (TUJ1) antibody. In control conditions, 38% of all explants did not grow any axon, compared to 19.5% for PTEN^−/−^ and less than 10% for PTEN^−/−^/SOCS3^−/−^+c-myc conditions (**Table 1**). This observation reflects in vivo conditions as WT optic nerves show very little regeneration compared to PTEN^−/−^ and PTEN^−/−^/SOCS3^−/−^+c-myc conditions. For the explants that grew more than one axon, we measured axon number and length in WT, PTEN^−/−^ and PTEN^−/−^/SOCS3^−/−^+c-myc conditions (**Figure 3G, H, I; Suppl Figure 2A, B, C, D**). To do so, we used the Scholl analysis-based plug-in Neurite-J(*18*) on Image J software. An automated method defined the explant borders and filtered out noise in the background, with manual correction if necessary. Subsequently, we used the automated Scholl analysis to measure the number of intersections (outgrowing neurites) at defined distances of the explants (50µm step) (**Suppl Figure 2E**). As expected, neurite outgrowth was very weak in control conditions, as only few axons came out of the explant and with a very short outgrowth (**Figure 3G, H, I; Suppl Figure 2**). PTEN^−/−^ explants showed an intermediate phenotype like in the in vivo condition. In contrast, for PTEN^−/−^/SOCS3^−/−^+c-myc explants, a very high number of axons grew far from the explant border (up to 8mm) (**Figure 3G, H, I; Suppl Figure 2**). Altogether, our experiments show that ex vivo adult retina explant cultures recapitulate perfectly in vivo growth phenotypes. Therefore, it is a relevant model to use to address the fine cellular and molecular mechanisms of axon regeneration.

### Laser guided axon ablation to mimic CNS lesion in a dish

As adult retina explant cultures recapitulate the in vivo phenotype of axon regeneration, we asked next how to mimic axon injury. Here, we wanted to address the precocious events as well as later ones at the level of a single axon (**Figure 4A**). We used laser ablation of single axons, in a similar approach to what is described in vivo in Drosophila or C. Elegans(*19, 20*) or in vitro in primary cultures of dissociated mouse embryonic neurons(*21, 22*). This technique allows the ablation of an axon without damaging the cell. Therefore, it enables to study early events that occurs after axon injury such as calcium dynamics or later events such as growth cone formation or guidance mechanisms (**Figure 1H**). For each experiment, we selected a single axon and followed it until its tip. We imaged the growth cone for 20min (1 image every two seconds) to ensure viability (**Video 1**). Because of the variability in axon length, we chose to define the lesion site about 100µm from the tip of the growth cone. Live imaging was performed up to 1 hour after the lesion (1 image every two seconds). In some cases, we observed that laser ablation was not efficient to cut the axon. Several scenarii are possible. In most cases, the laser is not aligned with the axon plane. Thus, the cut was effective by adjusting the focus. In other cases, laser power was not sufficient, depending on the thickness of the axon. We usually increased the attenuation plate (up to 64% transmission) and the laser power (up to 18%) to perform effective ablation (see **Material and Methods**). However, in order to avoid any unnecessary damage to the axon, we restrained the laser ablation to 2 attempts per axon. If the cut was incomplete or unsuccessful, we worked with another axon. For acquisitions that lasted over an hour, we studied one axon per explant.

**Figure 4:**
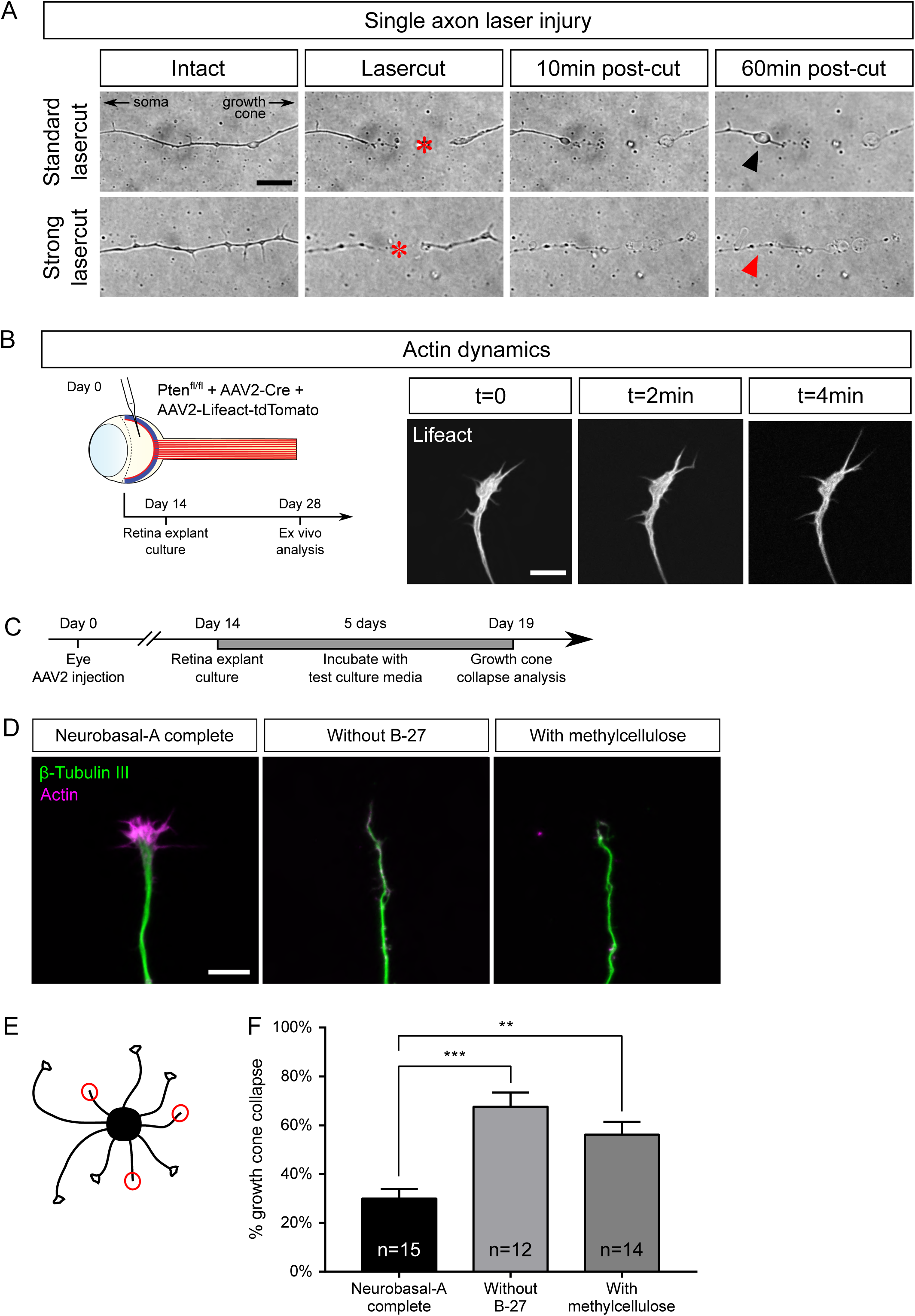
Examples of ex vivo analysis in adult retina explant cultures. (A) Single axon laser injury performed with MicroPoint Laser Illumination (Andor, Oxford Instrument). After lasercut (red star), a retraction bulb (black arrowhead) forms in the proximal part of the axon, while the distal part degenerates. In the case of a high-power (strong) lasercut, both proximal (red arrowhead) and distal parts degenerate. (B) Intravitreal injection of AAV2-Lifeact-tdTomato allows live visualization of actin cytoskeleton dynamics in a growth cone of adult retina explant culture. (C) Timeline of experiment of growth cone collapse assay in adult retina explant cultures. (D) Representative confocal pictures of growth cones of adult retina explant cultures in standard culture medium (Neurobasal-A complete), without B-27 and with methylcellulose. Scale bar: 10µm. (E) Schematic drawing showing the analysis of a collapse assay. Collapsed growth cones are indicated in red circles. (F) Quantification of growth cone collapse. Data expressed as means +/- s.e.m. Kruskal-Wallis test with comparison to control condition (Neurobasal-A complete). ** p<0.01, *** p<0.001.

Upon efficient laser ablation, the distal part of the axon undergoes degeneration, as described in vivo(*23*). Regarding the proximal part of the axon, there were two main outcomes after laser ablation: either a retraction or a fast degradation. In the first case, we observed axon retraction from the lesion site (**Figure 4A, Video 2**). Interestingly, this observation has also been reported in vivo(*24, 23*), especially in the case of spinal cord injuries. Thus, ex vivo adult retina explant cultures recapitulate the very first events following axonal lesion. In the second case, the degeneration spread fast all along the axon (**Figure 4A, Video 3**). In our ex vivo set up, this phenotype meant that the ablation power was too high and damaged the cell. Those axons were excluded from experiments.

### Adult retina explant cultures as useful tool to study molecular and cellular mechanisms of CNS regeneration: a few examples of application

Axon regeneration relies on several processes, such as growth cone formation, organelle movement or calcium dynamics. We used several experimental set-ups as proof of concept to show that adult retina explant cultures are the ideal tool to address these questions.

### * Growth cone dynamics

The growth cone is a highly motile structure at the end of growing axons. It is essentially composed of a central domain of microtubule and the periphery is formed by actin cytoskeleton(*25*). The growth cone is involved in axon elongation and guidance, as guidance receptors are expressed at the growth cone surface(*26, 27*). In the mature CNS, lesioned axons fail to form a growth cone, and growth cone formation is considered as a limiting step in CNS regeneration. Our recent work used this ex vivo set-up to show that DCLK2 (doublecortin like kinase 2), a structural protein regulating microtubules and actin, facilitates growth cone formation upon lesion and therefore induces axon regeneration in mature CNS(*28*). Here, we could study growth cone dynamics in live explants. To do so, we injected AAV2-Lifeact coupled with tdTomato into the vitreous body of PTEN^fl/fl^ mice together with AAV2-Cre (**Figure 4B**). Lifeact is a small probe that binds specifically to polymerized actin. Therefore, we could record growth cone dynamics as well as actin dynamics in live cultures (**Video 4**).

Growth cones could also be analyzed in fixed cultures. The growth cone acts as a target-tracking sensor of the navigating axon(*29*). Indeed, it expresses surface receptors that integrate environment signals such as guidance factors. When a growth cone comes across a repulsive cue, it will retract and change its direction of growth(*26*). It is well admitted that the early sign of repulsion is the collapse of the growth cone(*30*). This drastic change in the growth cone morphology is due to the depolymerization of the cytoskeleton, mainly actin(*27*). Adult retina explant cultures are very suitable to study mature axon response to environmental cues. Here we showed that growth cones elicit a specific collapse response that depends on culture medium composition. We tested several culture conditions, usually used for embryonic cultures, to address growth cone behavior. In particular, this allowed us to find the lowest baseline of growth cone collapse rate in adult retina explant cultures. We incubated explants either in complete (B27-containing) medium, or in medium without B27, or in medium supplemented with methylcellulose (**Figure 4C**). After 5 days, cultures were fixed and stained with anti-β Tubulin III (TUJ1) antibody and phalloidin, which labels polymerized F-actin (**Figure 4D**). For each explant, we counted the number of axons exhibiting a non-collapsed (spread) and collapsed (less than 2 filopodia and no lamellipodia) phenotype (**Figure 4E**). With the complete medium, the basal rate of growth cone collapse was about 30%, which is slightly higher than embryonic cultures of cortical neurons (around 20%(*31*)). In contrast, unlike young neurons(*32*), mature RGC axons are very sensitive to B27 deprivation or methylcellulose supplementation. In both conditions, we observed an increase of the basal rate of growth cone collapse (more than 60%) (**Figure 4F**). We concluded that it is essential to supplement the medium with B27 to minimize growth cone collapse rate in mature cultures. These cultures are different from embryonic neurons and do not respond equally to media composition. In addition, this experiment showed that adult axons are able to respond to their environment ex vivo. This opens up the possibility to use adult retina explant cultures to decipher how adult axons integrate environmental signals, and in particular guidance cues.

### * Axonal transport

Axons are busy highways for many types of transport. The transport of specific organelles, such as mitochondria, has been shown to be critical to achieve regeneration(*33*). High frequency moving mitochondria have been linked with axons showing strong growth capability(*34*). However, studying mitochondria dynamics in vivo remains technically challenging(*35*). Here we show that adult retina explant cultures are a good tool to address this question. We used two approaches to label mitochondria in live axons: viral infection and incubation with a specific dye. We infected PTEN^fl/fl^ adult RGC with AAV2-Cre together with AAV2-MitoDsRed, which labels all mitochondria with DsRed (**Figure 5A**). In this experimental design, all cells of the explant expressed MitoDsRed, which could be tracked with live confocal imaging (**Video 5**). The second approach is based on the use of live tracking cell-permeant dyes more adapted to study different organelles from the same culture. In this case, we put in culture adult retina explants from PTEN^fl/fl^ mice, whose RGC were infected with AAV2-Cre. After two weeks, explants were incubated for 5-10 min with MitoTracker, a specific mitochondrion-selective dye that accumulates in active mitochondria; or for 20-30min with LysoTracker, which labels acidic organelles and enables lysosome tracking. We imaged organelle dynamics 5min before and 5min after laser guided axon lesion (**Figure 5B, C**) and performed a kymograph analysis in the proximal region of the axon close to the lesion site (about 15µm). Interestingly, we found that in mature intact axons, most mitochondria (90%) are stationary (**Figure 5D-J**), similar to what was shown in vivo(*33, 36*). There was only 10% of slowly moving mitochondria and the laser guided lesion did not affect this proportion (**Figure 5I, Video 6A, B**). Anterograde and retrograde movements were equally not affected by axon injury (**Figure 5G**). In contrast, we observed that lysosomes, tracked with LysoTracker, were extremely motile (**Figure 5K-Q, Video 7A, B**), with a bidirectional movement predominantly retrograde, as described for maturing lysosomes tracked with LysoTracker in axons(*37*). Indeed, 90% of lysosomes were moving (both anterogradely and retrogradely) in intact axons (**Figure 5P**) with an average retrograde speed of 0.6µm/s and an average anterograde speed of 0.3µm/s (**Figure 5N**). After axon ablation, only 30% of lysosomes kept moving (**Figure 5N-P**), with an average speed reduced to less than 0.06µm/s in both directions (**Figure 5N**). The pausing time increased from 50% in intact conditions to almost 100% after axon injury (**Figure 5Q, Video 7A, B**). Therefore, axon ablation disrupts lysosome movements in adult axons.

**Figure 5:**
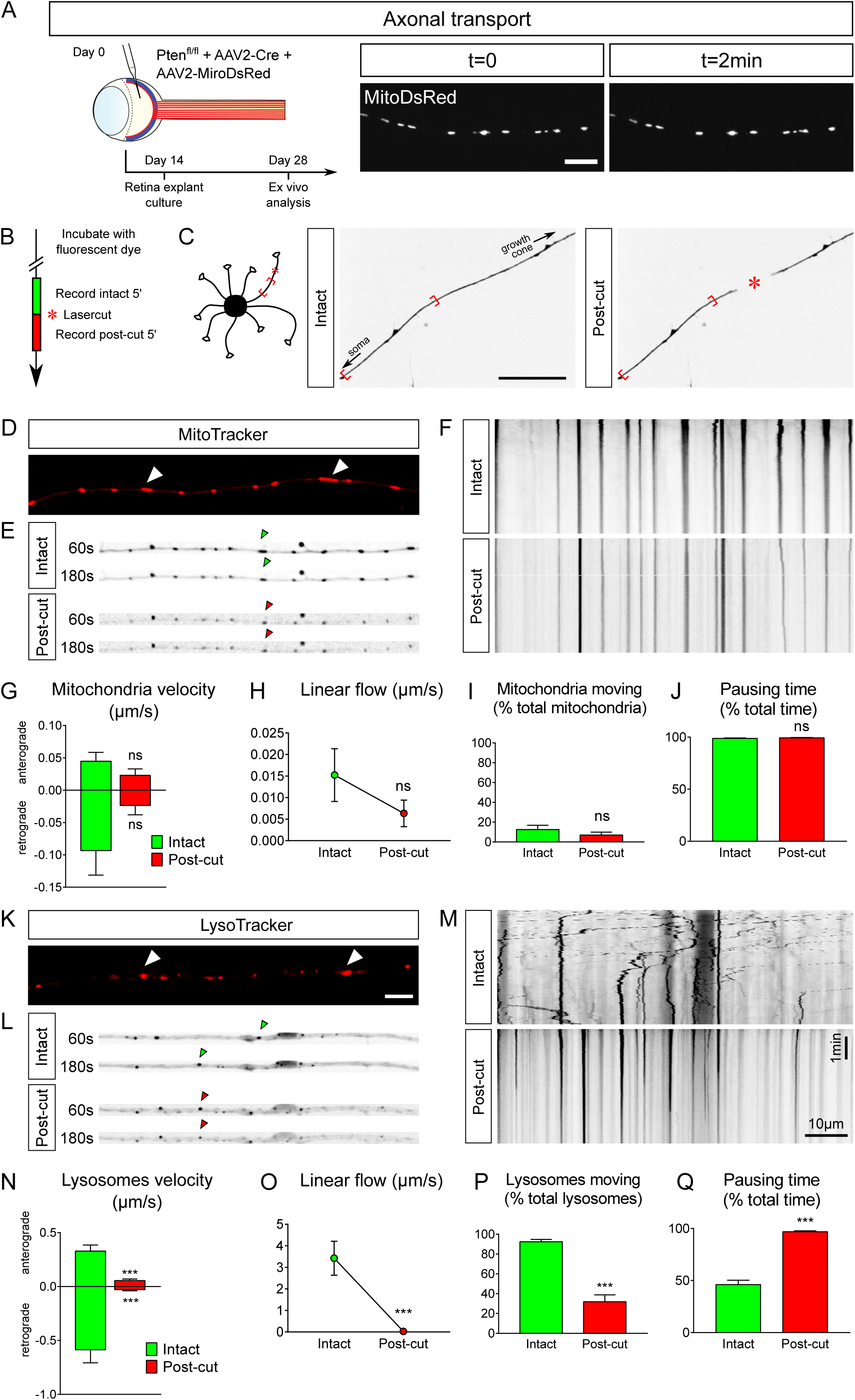
Ex vivo analysis of axonal transport in intact and injured conditions in adult retina explant cultures. (A) Intravitreal injection of AAV2-MitoDsRed in the eye allows live visualization of mitochondria axonal transport in adult retina explant culture. Scale bar: 10µm. (B) Timeline of live analysis. Adult retina explant cultures are incubated with fluorescent dyes, which allows live tracking of lysosomes or mitochondria. (C) Representative pictures of a single axon before and after laser injury (intact and post-cut). The red star indicates the site of laser injury. The blue brackets indicate the region recorded for axonal transport analysis. Scale bar: 50µm. (D) Representative confocal picture of MitoTracker Red labeling of an axon, showing mitochondria (white arrowheads). Scale bar: 5µm. (E) Representative pictures of mitochondria transport in an axon in intact and post-cut conditions. Arrowheads show tracking of a single mitochondria over time. (F) Representative kymographs of mitochondria tracking in intact and post-cut conditions. (G-J) Quantifications of mitochondria velocity, global linear flow, number of moving mitocondria and pausing time. n=9 axons. Mann-Whitney tests. ns: not significant. (K) Representative confocal picture of LysoTracker Red labeling of an axon, showing lysosomes (white arrowheads). Scale bar: 5µm. (L) Representative pictures of lysosme transport in an axon in intact and post-cut conditions. Arrowheads show tracking of a single lysosome over time. (M) Representative kymographs of lysosome tracking in intact and post-cut conditions. (N-Q) Quantifications of lysosome velocity, global linear flow, number of moving lysosomes and pausing time. n=9 axons. Mann-Whitney tests. *** p<0.001. (G-J) and (N-Q) Data expressed as means +/- s.e.m.

Altogether, our experiments prove that adult retina explant cultures enable to sustain adult axons in culture and to isolate various features critical to achieve axon regeneration in the mature CNS.

## DISCUSSION

Understanding CNS regeneration has been a major challenge for centuries since Ramon y Cajal’s first observations that unlike the PNS, CNS neurons fail to form a growth cone to achieve successful regrowth after lesion(*38*). One of the main issues is to decipher the mechanisms underlying mature axon growth. Indeed, it is challenging to sustain adult CNS neurons in culture. Therefore, most studies address the regulation of axon regeneration by using embryonic cultures as an in vitro tool. However, this approach presents a major caveat: developing neurons have the intrinsic ability to grow their axon after lesion. The molecular and cellular growth pathways may not be the same during development and regeneration. This explains partially why several pathways characterized in vitro using such embryonic cultures do not induce extended regeneration in vivo(*7, 39–43*). To circumvent this discrepancy, we propose to use adult retina cultures. Retina explants are relevant because the visual system is part of the CNS and the optic nerve presents the same features as the rest of the CNS regarding injury: RGC axons fail to regenerate and to survive after lesion. Using the optic nerve as a model, the modulation of neuronal intrinsic capabilities has been shown to be key to promote axon regeneration in the mature CNS. Consequently, over the past decade, the optic nerve has become a gold-standard model to address axon regeneration in the CNS. Importantly, axon regeneration is triggered in a similar way in RGC and in other CNS neurons, such as cortico-spinal neurons or dopaminergic neurons, upon activation of mTOR(*9, 44*).

Here, we set up adult retina explant cultures to address the molecular and cellular mechanisms underlying axon regeneration in mature CNS neurons. Explants from PTEN^fl/fl^/YFP-17 retina allowed us to show that the vast majority of outgrowing neurites are RGC axons. Thus, the neuronal population of interest (RGC) are able to grow axons in our ex vivo set-up. Second, adult retina explant cultures enable to decipher regeneration mechanisms at the level of single axons. Third, the timeline of experiments in vivo and ex vivo is exactly the same, so neurons are studied exactly at the same stage in vivo and ex vivo. In addition, we showed that the retina explant cultures display the same extent of growth as what is observed in vivo. This result suggests that the functional mechanisms underlying axon regeneration are maintained in the ex vivo system. For these reasons, our ex vivo experimental model recapitulates very precisely the molecular and cellular events that control axon growth in the adult CNS.

The clearing approach gave us a full phenotypic view of axon regeneration from the eye to the brain with no bias. In parallel, adult retina explant cultures allowed us to address the fine molecular and cellular events that control axon regeneration. Understanding those mechanisms is crucial to design translational approaches and propose a treatment for patients suffering from CNS traumatic injuries, but also from neurodegenerative disorders, which raise similar questions of neuroprotection and neuroregeneration in the adult CNS. It is interesting to note that WT adult retina explants display very little neurite outgrowth, which corresponds exactly to the failure of regeneration in the mature WT CNS. Therefore, axonal growth in WT adult retina explants can be used as a baseline to test pro-regenerative molecules in a pharmacological approach of CNS regeneration. Thereafter we give a non-exhaustive overview of the experimental applications that adult retina explant cultures offer.

While axons from the PNS are able to form a growth cone within hours after lesion(*45*), CNS axons fail to achieve this step and instead, the tip of the lesioned axon is sealed by a retraction bulb(*23*). It is believed that this abortive attempt to form a functional growth cone is part of the mechanism underlying the failure of the mature CNS to regenerate and is the limiting step to reach axon growth(*27, 46, 47*). Therefore, it is crucial to identify molecular and cellular regulators of growth cone formation in CNS axons. In this context, adult retina explant cultures are highly handful to address this question. To study growth cone reformation after axonal lesion, we combined live imaging and laser ablation of a single axon. In a previous study we showed that the structural protein DCLK2 (doublecortine like kinase 2) enhances axon regeneration by inducing growth cone formation through actin cytoskeleton stabilization(*28*). This set-up allows to analyze regenerating growth cones in live and in detail, and thus to study how mature axon growth cones respond to their environment. Furthermore, the behavior of growing axons towards their target is still elusive. While modulation of neuronal intrinsic abilities to promote axon regrowth has led to long distance regeneration, the formation of functional circuits still remains challenging. Therefore, deciphering the interaction between regenerating axons, and especially their growth cones, and molecules expressed in the environment will be key to drive these axons to relevant, functional targets.

Moreover, even if molecular pathways that control axon regeneration have been uncovered, the underlying mechanisms are still difficult to elucidate. For example, discrepancy in axonal transport has been linked to neurodegenerative diseases in many studies but remains to be fully characterized(*48, 49*). Accelerating mitochondria transport leads to axon regeneration by supplying healthy mitochondria and rescuing energy deficits in the injured axon(*33*). Overexpression of enhancers of mitochondria movement, such as the protein Armcx1, triggers axon regeneration in the optic nerve(*50*). In our ex vivo system, mitochondria could be labelled using viruses that express MitoDsRed or using a specific tracking dye added extemporary into the culture media. Here we showed that in mature intact axons mitochondria are stationary, as observed in vivo for the majority of mitochondria(*51, 36*), although intravital imaging of mitochondrial axonal transport in mouse RCG shows some active transport near the soma in intact conditions(*35*) that we did not studied ex vivo as we analyzed distal regions of the axon. In contrast to adult axons, mitochondria are motile in developing neurons(*52, 53*). This opposite behavior in developing and mature axons could create a bias when it comes to study the contribution of mitochondria during regeneration, again showing the importance of an ex vivo model as close to in vivo conditions as possible. In adult axons, we observed that mitochondria are stationary in intact conditions and following the lesion. It would be interesting to test pathways known to regulate their motility and recruitment and address the outcome upon lesion(*33, 34, 54, 55*).

Interestingly, other organelles such as lysosomes show a high dynamics in mature axons, with a tendency for retrograde movement that correlates with their maturation (*37*). Upon lesion we observed a significant decrease of their velocity. Yet, it is now understood that correct lysosome polarized movement in the axon is essential for maintaining global axon functioning, including growth cone dynamics(*56*). More generally, lysosome transport deficits are associated with many neurodegenerative disorders(*57–59*). Thus, the almost complete interruption of lysosome dynamics caused by axonal injury may strongly impair the regenerative capacity of adult CNS axons. It has been previously shown that axon lesion induces the disruption of actin and microtubule cytoskeleton(*27*). Most of the organelles use the microtubules as highways to navigate within the cells(*60*). Therefore, it is not surprising that lysosome movements drop upon lesion as microtubules depolymerize. It would be interesting to address organelle movement when the cytoskeleton is preserved upon taxol treatment, which stabilizes microtubules, or upon overexpression of microtubule-associated and actin regulating proteins such as DCLK2.

In this study we showed that adult retina explant cultures are the ideal ex vivo system to explore the molecular and cellular events that occur upon axon lesion in the mature CNS. We described a non-exhaustive series of experimental applications of this model, such as characterization of growth cone behavior and study of organelle transport in the axon. This model recapitulates the in vivo phenotype and offers to characterize finely adult CNS axons at a single axon level, in a biological set-up far more relevant and accurate to the field of CNS repair than embryonic neuronal cultures. It is critical to understand these events to find new targets to achieve repair of the mature nervous system and formation of functional circuits. We expect that this ex vivo system will shed light on novel cellular and molecular mechanisms underlying axon regeneration, leading to the development of effective therapeutic strategies for CNS repair. Our model will also be extremely useful to address neurodegenerative diseases that raise the same unanswered questions of neuroprotection and neurodegeneration in the adult CNS.

## METHODS

### Animals

Animal care and procedures were performed according to the Grenoble Institute of Neuroscience, French and European guidelines.

We used PTEN^fl/fl^/YFP-17, PTEN^fl/fl^ and PTEN^fl/fl^/SOCS3^fl/fl^ mice lines in this study, regardless of their sex, aged at least 4 weeks.

### Intravitreal virus injection

4-weeks-old animals were anesthetized with Ketamine (60-100mg/Kg) and Xylazine (5-10mg/Kg). We used the same protocol as described before to perform intravitreal injections(*5*). The external edge of the eye was clamped using an artery clamp to display the conjunctiva. Using a glass micropipette connected to a Hamilton syringe, 1µL of Adeno-associated type 2 viruses (AAV2-Cre, AAV2-CNTF, AAV2-c-myc, AAV2-mitodsRed or AAV2-Lifeact-tdTomato; at least 10^11 viral particles per ml) were injected into the vitreous body of the eye. Mice with eye inflammation or damage were excluded from the rest of the experiments. Following the same procedure, 1µL of CTB-555 (Invitrogen) at 1µg/µL was injected into the vitreous body of the eye 2 days before termination.

### Optic nerve crush

Two weeks after viral injection, animals used to address axon regeneration underwent optic nerve crush. After anesthetizing the animals with Ketamine (60-100mg/Kg) and Xylazine (5-10mg/Kg) we opened the conjunctiva with fine scissors. Carefully we slided dilating forceps (Fine Science Tools) in-between the two arteries behind the eye ball to expose the optic nerve underneath. Then the optic nerve was pinched for 5s using jeweler’s forceps (Dumont #5 forceps-Fine science tools) 1-2mm behind the eye ball. Animals with unstoppable heavy bleeding were excluded from the study.

### Retina explant culture

The procedure is described in **Figure 1**. Two weeks after virus injection, animals were sacrificed by cervical dislocation following the institution’s guidelines. Eyes were removed quickly using Dumont’s forceps #5 and put in ice-cold Hibernate A medium without calcium and magnesium (Invitrogen). Under a dissection scope, the eyeball was opened and cut along the line between the anterior and posterior chambers of the eye using Spring scissors (Fine Science Tools). The retina, attached in the inferior part of the eye ball, was dissected out using #5 forceps and placed in a new dish containing ice-cold Hibernate A (Gibco) medium without calcium and magnesium. Using a scalpel blade, the retina was cut into small pieces (about 500µm in diameter). Glass coverslips or glass bottom dishes (MatTek) were first coated overnight at room temperature with Poly-L-Lysine (0.5mg/mL-Sigma Aldrich) and for at least 2 hours with Laminin (20ug/mL-Sigma Aldrich). After several washes in water, glass coverslips or glass bottom dishes were coated with a thin layer of coating media (Hibernate A (Gibco), 0,4% Methylcellulose (Sigma-Aldrich), 2% B27 (Gibco), 20mM L-Glutamine (Gibco)). Each retina explant presents a slight curvature. The layer of RGC is on the concave side of the explant. This side was placed on the coated glass coverslip or glass bottom dish. We put 2-4 explants per dish. After 5 minutes at room temperature, culture medium (Neurobasal-A (Gibco), 2% B27 (Gibco), 20mM L Glutamine (Gibco) and 5000 Units of Penicillin/Streptomycin (Gibco)) warmed at room temperature was gently added in the dish. At this step, all explants should be attached on the coverslip or glass bottom dish, otherwise they detach and should be removed from the culture. Cultures were grown at 37°C with 5% CO_2_ for 2 weeks.

### Immunostaining

For explant cultures, samples were fixed with 8% PFA and 3% sucrose diluted 1:1 directly in culture media. After several washes, cultures were permeabilized at room temperature for 10min in Triton X-100 (Sigma Aldrich) 0.1% in PBS. Then samples were incubated with primary antibodies for 2h at room temperature (anti-β Tubulin III (TUJ1-Covance 1:400), anti-Tau (Millipore, 1:250), anti GFP (AbCAM, 1:500)) in blocking solution (BSA (Sigma Aldrich) 3% in PBS). Finally, after several washes in PBS explants were incubated at room temperature for 1 hour in secondary antibodies diluted in the blocking solution (at 1:500 for Alexa-coupled antibodies (Life Sciences)) and 1:400 for TRITC-conjugated phalloidin (Sigma-Aldrich)) and then mounted using Fluoromount-G (Southern Biotech).

For retina sections, mice were intracardially perfused with ice-cold PFA. After dissecting out the eye balls and the optic nerves, samples were post fixed overnight at 4°C in 4% PFA. Eye balls and optic nerves were then separated. Eye balls were dehydrated in 15% sucrose for at least 48h at 4°C. After cryosectionning (14µm, Leica), samples were kept at −20°C. For immunostaining, slides were defrosted at room temperature for 20min. After several washes in PBS, samples were incubated for 1hour at room temperature in blocking solution (BSA (Sigma Aldrich) 3%, Triton X-100 (Sigma Aldrich) 0,5% in PBS). Samples were incubated with primary antibodies overnight at 4°C (anti-RBPMS (Millipore, 1:250), anti-GFP (AbCAM, 1:500)). Finally, slides were incubated at room temperature for 2 hours in secondary antibodies diluted in the blocking solution (at 1:500 for Alexa-coupled antibodies (Life Sciences)) and then mounted using Hoechst-containing Fluoromount-G (Southern Biotech).

### Whole optic nerve clarification

Clarification procedure is adapted from(*61*). After several washes in PBS, fixed optic nerves were progressively dehydrated in Ethanol (50%, 80%, 95 and 100%), then incubated in Ethanol 100% overnight at 4°C. The next day, samples were incubated for 2hours at room temperature in Hexane (Sigma Aldrich). Samples were then transferred in a Benzyl Benzoate/Benzyl Alcohol solution (Sigma Aldrich, 2:1) and stored in dark at 4°C until imaging.

### Imaging

For whole optic nerve imaging, we used the DragonFly spinning disc confocal from Andor. We took z stacks (2µm for each z step) to scan the entire width of the cleared optic nerves. Then we stitched images (with at least 10% of overlap) using a custom stitching module in Metamorph. We used the maximum z projection (performed with Metamorph) to visualize and quantify the extent of regeneration.

### Live imaging and laser ablation

All live imaging and laser ablation experiments were performed with PTEN^−/−^ retina explant cultures or PTEN^−/−^/Thy1-YFP retina explant cultures. For fluorescence live imaging, culture medium was replaced with unsupplemented Hibernate-A with no phenol red (BrainBits). Axons or growth cones were imaged with the DragonFly spinning disc confocal microscope from Andor, with 1 image per second. For standard DIC imaging and laser ablation, retina explant cultures were left to equilibrate in the humidified chamber at 37°C and 5% CO_2_ for 15min. We chose axons that were isolated from their neighbors to avoid working with fasciculate axons and we checked axon health by observing and recording growth cone dynamics for 20min before starting the experiment. Laser ablation was performed with Micropoint (Andor Technologies) and controlled with Metamorph imaging software. The galvo positions of the Micropoint were calibrated before each experiment to ensure accurate targeting of the axon. The cut was monitored visually with DIC illumination and recording. Laser ablation settings were: laser power set to 10%, with possible increase up to 18%; number of pulses set to 4; attenuation plate set to 25% transmission, with possible increase to up to 64%. No more than 2 attempts of laser ablation were made to avoid rapid and irreversible degeneration of the axon. Growth cones and/or axons were recorded for 20min before laser ablation, then 1hour after laser ablation, with 1 image every two seconds. Laser ablation was performed either close to the growth cone (about 100µm), or anywhere in the axon between the explant and the growth cone for organelle tracking, in a region where the axon was straight enough to maximize laser ablation efficiency. For organelle tracking, culture medium was replaced with unsupplemented Hibernate-A with no phenol red (BrainBits). Retina explant cultures were incubated with a live tracking dye, either MitoTracker (ThermoFisher, 0.1µM final concentration) for 5min, or with LysoTracker (ThermoFisher, 0.1µM final concentration) for 30min. Fluorescence was recorded in single axon for 5min before laser ablation, then for 5min after laser ablation, with 1 image every second. Up to 3 axons of the same explant could be recorded.

### Axon regeneration quantification

Quantifications were based on the maximum projection of the z-stack acquisition of transparent optic nerves (16-bit images). Using ImageJ, the injury site was manually defined with a straight line as the site where CTB labelling produces a “step” of intensity in the optic nerve (from saturated signal of intact fibers in the proximal part to lower signal of regenerating fibers in the distal part). The fluorescence intensity profile was measured at specific distances from the injury site (200, 500, 750, 1000, 1500, 2000, 2500 and 3000µm) along a line manually drawn orthogonally to the optic nerve, and with a length corresponding to the optic nerve width. The intensity profile was also recorded in a region with no regeneration (background measurement). All these steps were automatized using ImageJ. Integrated fluorescence intensity was calculated at each step using R and normalized to the optic nerve width at the same step (that may vary along the optic nerve). The integrated intensity was normalized to the maximal intensity value of all steps in the regenerating region to account for variations between optic nerves. Finally, the normalized integrated intensity of background was subtracted from the normalized integrated intensity at each step, and the results were plotted in arbitrary units as a function of the distance from the injury site.

### Explant axon outgrowth quantification

Explants were imaged with epifluorescence microscopy with automatic stitching (Axioscan slide scanner Scan.Z1, Zeiss). Explants with no or little axon outgrowth (0 or 1 axon) were removed and analyzed separately (**Table 1**). For the rest of the explants, axon outgrowth was quantified with a Scholl analysis using the ImageJ plug-in Neurite-J(*18*). Definition of the explant and background noise filtering were automatically performed with manual correction if necessary, as described by the plug-in developer. The number of neurites intersects was determined by the Scholl analysis with a step of 50µm. Data were plotted as the number of neurite intersects as a function of the distance to the explant border.

### Organelle live tracking quantification

The Image-J plug-in KymoToolBox(*62*) was used to quantify organelle dynamics (mitochondria or lysosomes). The region of interest designed as a segmented line was defined along the axon in the proximal part, about 15µm from the laser ablation point and of about 120µm in length (**Figure 5C**). Kymographs were automatically drawn using the Draw Kymo command. A total of 9 axons were analyzed for each organelle. For kymograph analysis, 9 to 18 lysosomes and 13 to 25 mitochondria were selected and the trajectories manually drawn with segmented lines. Kymographs were analyzed with the Analyse Kymo command with the following parameters: minimum speed = 0.02µm/s. Several kinetics parameters were calculated according to(*63*), as the following: anterograde velocity = Vma (µm/s) = Anterograde Distance (µm) / Time (s), retrograde velocity = Vmr (µm/s) = Retrograde Distance (µm) / Time (s), pausing time = average(pausing time per axon), linear flow rate = Q (µm/s) = |Vma| * Number of anterograde organelles + |Vmr| * Number of retrograde organelles.

### Statistical analysis

Statistical analysis was performed with GraphPad Prism (version 7.00). For each dataset, the Shapiro-Wilk test was used to assess normal distribution (P-value >= 0.01). Datasets normally distributed were analyzed with an unpaired Student’s t-test (for comparison of two conditions) or an ANOVA test (for multiple comparisons). Datasets that were not normally distributed were analyzed with a Mann-Whitney test (for comparison of two conditions) or a Kruskal-Wallis test (for multiple comparisons).

## Supporting information

Growth cone of adult retina axon

standard laser cut of single axon

strong laser cut of single axon

actin dynamics in growth cone

mitochondria in intact axon

mitochondria in intact axon

mitochondria in injured axon

lysosomes in intact axon

lysosomes in injured axon

## AUTHOR CONTRIBUTIONS

Conceptualization H.N and S.B; Methodology HN and SB. Experiments were performed by JS and CT. Supervision HN and SB. Original draft and editing HN SB JS and CT. Mice colony handling and technical assistance FA. Funding acquisition HN and SB.

## ACKNOWLEDGEMENTS

We would like to thank Prof Zhigang He for critical reading of this manuscript and Dr. Chen Wang for technical assistance (Boston Childrens Hosptial-Harvard Medical School). We thank Charlotte Corrao and Mohamed-Elmehdi Boughanmi for virus preparation. This work was supported by a grant from ANR to HN (C7H-ANR16C49) and SB (ANR-18-CE16-0007), from European Research Council (ERC-St17-759089) and NRJ Foundation to HN. This work was supported by grants from the French National Research Agency in the framework of the “investissements d’Avenir” program (ANR-15-IDEX-02 NeuroCoG (HN, SB and CT)). JS is supported by UNADEV/AVISAN grant (Appel à projets 2017 “Maladies de la vision, origine et traitement”) and by Fondation pour la Recherche Médicale (FRM) postdoctoral fellowship (SPF201909009106). This work was supported by the Photonic Imaging Center of Grenoble Institute Neuroscience (Univ Grenoble Alpes – Inserm U1216) which is part of the ISdV core facility and certified by the IBiSA label.

## FIGURE LEGENDS

**Supplementary Figure 1:**
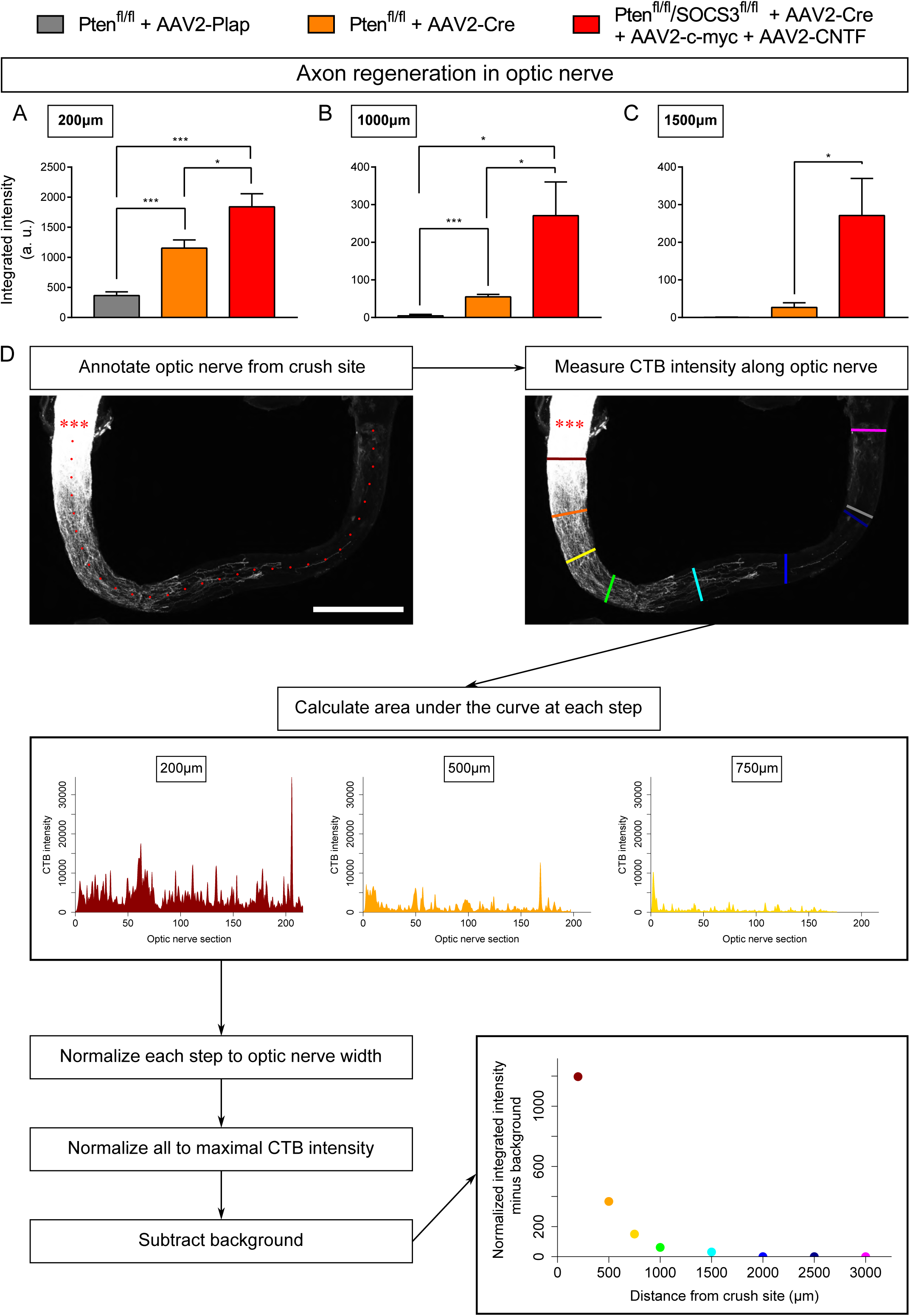
Quantification of axon regeneration in optic nerves whole-mount. (A-C) Comparison of optic nerves from control (PTEN^fl/fl^ + AAV2-Plap), PTEN-deleted (PTEN^fl/fl^ + AAV2-Cre) and PTEN/SOCS3 co-deleted c-myc-overexpressing (PTEN^fl/fl^/SOCS3^fl/fl^ + AAV2-Cre + AAV2-CNTF + AAV2-c-myc) conditions, at different distances from the injury site. Data are expressed as means +/- s.e.m. Unpaired t-tests. * p<0.05, ** p<0.01, *** p<0.001, **** p< 0.0001. (D) Principle of axon regeneration analysis in whole transparent optic nerves. Representative confocal picture of CTB-labelled optic nerve whole-mount from a PTEN^fl/fl^ mouse with prior intravitreal injection of AAV2-Cre. Red stars indicate the injury site. The confocal picture is annotated with ticks spaced 100µm along the optic nerve. CTB-555 intensity is measured at defined steps (coloured lines) of optic nerve, with background measurement in a region with no axon regeneration (grey line). At each defined step, the area under the curve is measured and normalised to optic nerve width. Values are normalised to the maximal intensity value of all defined steps. Finally, the background value is subtracted. Scale bar: 500µm.

**Supplementary Figure 2:**
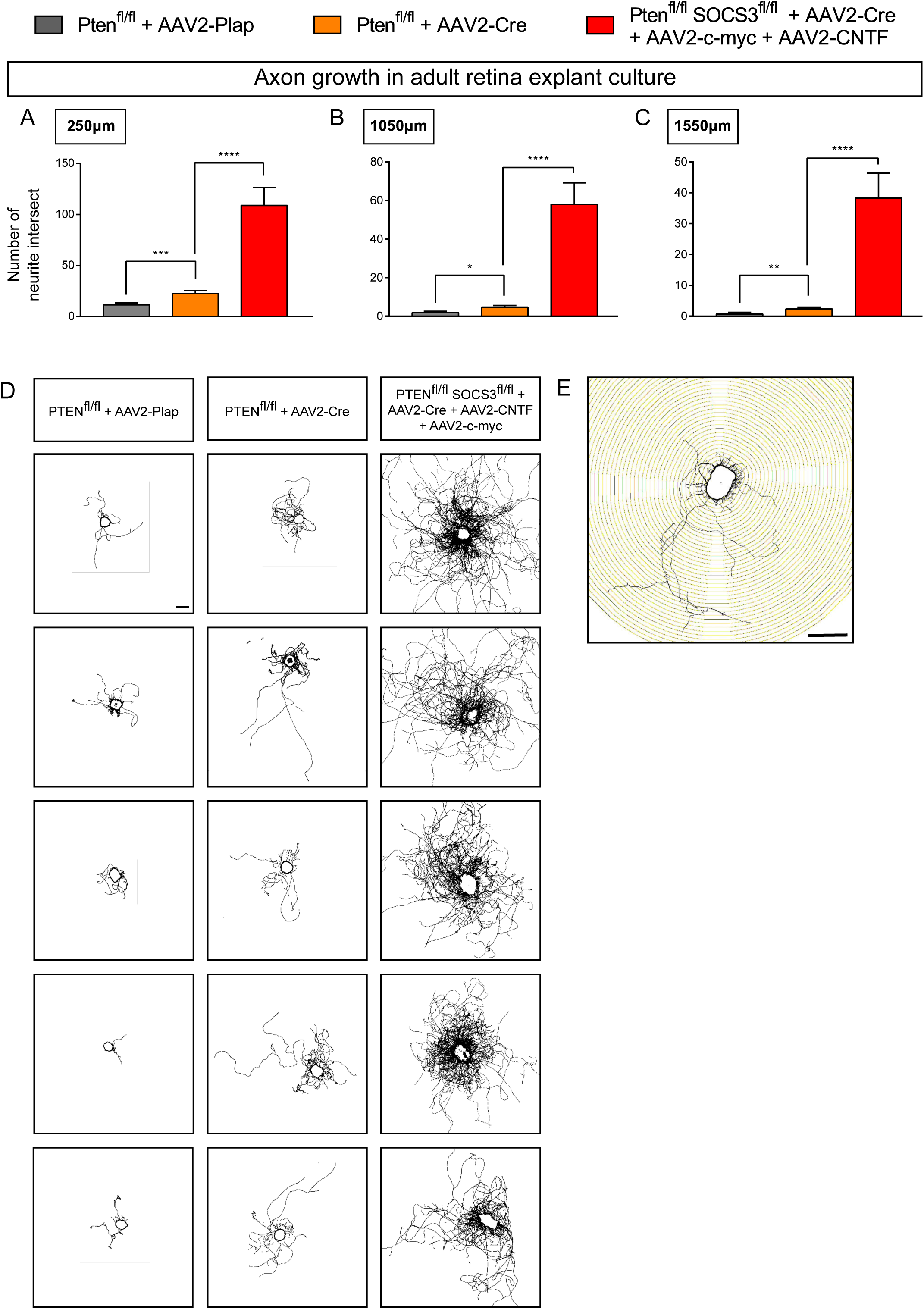
Quantification of axon growth in adult retina explant cultures. (A-C) Comparison of explant cultures from control (PTEN^fl/fl^ + AAV2-Plap), PTEN-deleted (PTEN^fl/fl^ + AAV2-Cre) and PTEN/SOCS3 co-deleted + c-myc-overexpressing (PTEN^fl/fl^/SOCS3^fl/fl^ + AAV2-Cre + AAV2-CNTF + AAV2-c-myc) conditions, at different distances from the explant. Data are expressed as means +/- s.e.m. Unpaired t-tests. * p<0.05, ** p<0.01, *** p<0.001, **** p< 0.0001. (D) Representative pictures of adult retina explants after 2 weeks in culture, from control (PTEN^fl/fl^ + AAV2-Plap), PTEN-deleted (PTEN^fl/fl^ + AAV2-Cre) and PTEN/SOCS3 co-deleted + c-myc-overexpressing (PTEN^fl/fl^/SOCS3^fl/fl^ + AAV2-Cre + AAV2-CNTF + AAV2-c-myc) conditions. Axons are labelled with anti-β Tubulin III (TUJ1) antibody. Scale bar: 500µm. (E) Example of Scholl analysis for quantification of number of axon intersect.

